# Differential isoform-specific control of KCC2 function in developing cortical neurons

**DOI:** 10.64898/2026.01.25.701648

**Authors:** Carla Pagan, Pauline Weinzettl, Marika A Markkanen, Marion Russeau, Matti S Airaksinen, Jean Christophe Poncer

**Author notes:** Corresponding author Jean Christophe Poncer Paris Brain Institute Hôpital Pitié-Salpêtrière 47 boulevard de l’Hôpital 75013 Paris, France.

## Abstract

The K-Cl cotransporter KCC2 is essential for fast synaptic inhibition in the mature brain. It is encoded by a single gene and expressed as two isoforms: KCC2a and KCC2b, which differ in their N-terminal domains. While KCC2b is predominant, the function of the weakly expressed KCC2a isoform remains unclear. Here, we reveal that KCC2a is a potent, bidirectional regulator of KCC2b membrane stability and function in cortical neurons. In immature neurons, where WNK-SPAK kinase activity is high, KCC2a promotes SPAK-dependent phosphorylation of KCC2b at Thr1007, which likely contributes to hindering its membrane expression and function. Conversely, in mature neurons with low basal WNK-SPAK activity, KCC2a promotes KCC2b expression, clustering, and function. At this stage, although accounting for less than 5% of total KCC2 mRNA, KCC2a is enriched in dendrites and within KCC2 clusters, where it prevents clathrin-mediated KCC2b endocytosis, as well as polyubiquitination and proteasomal degradation. Thus, KCC2a acts as a developmental switch that first inhibits KCC2b during early development and then ensures its membrane stability to support effective synaptic inhibition in the adult brain.

## Introduction

The KCC2 neuronal K-Cl cotransporter, which is encoded by the *SLC12A5* gene, is essential for maintaining the low intracellular chloride concentration required for fast hyperpolarizing synaptic responses mediated by inhibitory neurotransmitters, such as gamma-aminobutyric acid (GABA) and glycine, in the mature central nervous system. Upregulation of KCC2 expression is a hallmark of neuronal maturation (Rivera *et al*, 1999; Virtanen *et al*, 2021), mediating the developmental shift in GABAergic signaling from depolarizing to hyperpolarizing (Ben-Ari *et al*, 2002). Due to its pivotal role in chloride homeostasis and GABA signaling, KCC2 dysfunction has been associated with many neurological and psychiatric disorders (Kaila *et al*, 2014), including epilepsy (Huberfeld *et al*, 2007), neuropathic pain (Coull *et al*, 2003), and Alzheimer’s disease (Keramidis *et al*, 2023).

The *SLC12A5* gene generates two full-length protein isoforms, KCC2a and KCC2b, via alternative promoters and splicing (Uvarov *et al*, 2007). These isoforms only differ in their unique N-terminal sequences. Whereas the fundamental role of KCC2b in chloride transport is well-established and primarily drives the developmental switch in GABA signaling in cortical neurons (Uvarov *et al*, 2009), the regulatory profile and physiological function of KCC2a remain elusive.

Genetic studies suggest a critical and transient role of the KCC2a isoform, particularly in the early stages of postnatal development. Complete genetic ablation of *Slc12a5* in mice results in death at birth due to severe motor deficits (Hubner *et al*, 2001). However, mice lacking only KCC2b (while retaining KCC2a) survive for 2–3 weeks, suggesting that KCC2a expression alone is sufficient to support vital functions during the immediate postnatal period (Woo *et al*, 2002). Conversely, KCC2a-deficient mice exhibit transient respiratory anomalies at birth, including low breathing rates and apneas, resulting from abnormally strong inhibition from the pons (Dubois *et al*, 2018). This functional divergence is mirrored by the distinct developmental expression patterns of KCC2 isoforms. KCC2a accounts for a substantial fraction (50-65%) of total KCC2 RNA in the cortex, brainstem, and spinal cord on embryonic day 17 (Uvarov *et al*, 2007; Uvarov *et al*, 2009). However, its relative contribution decreases drastically in the adult forebrain, where KCC2b becomes the predominant isoform (>90%) (Markkanen *et al*, 2014; Uvarov *et al*, 2007). Therefore, the differential regulation and developmental switch of KCC2 isoforms appear critical for circuit function, especially during the perinatal period.

Both KCC2a and KCC2b were shown to function as K-Cl cotransporters (Uvarov *et al*, 2007; Uvarov *et al*, 2009) and interact to form heterodimers in both heterologous cells and rat neurons (Uvarov *et al*, 2009). However, their distinct N-terminal sequences dictate different regulatory mechanisms, enabling the fine-tuning of K-Cl transport activity. In particular, the N-terminus of KCC2a, but not KCC2b, contains a conserved (RFxV/I) binding motif for the Ste20-related proline-alanine-rich kinase (SPAK) (Uvarov *et al*, 2009), an effector of the WNK (with-no-lysin) kinase pathway known to regulate cation-chloride cotransporters (de Los Heros *et al*, 2014; Friedel *et al*, 2015; Heubl *et al*, 2017). Activation of the WNK-SPAK pathway results in the phosphorylation of KCC2b at Thr1007, leading to a subsequent decrease in membrane stability and chloride extrusion function (Friedel *et al*, 2015; Heubl *et al*, 2017). In heterologous cells, SPAK was found to co-immunoprecipitate specifically with recombinant KCC2a, but not with KCC2b (Markkanen *et al*, 2017). Functionally, the overexpression of SPAK significantly decreased the transport activity of KCC2a, yet it had no inhibitory effect on KCC2b in the absence of KCC2a (Markkanen *et al*, 2017). These results suggest that KCC2a may be essential for the inhibitory regulation of KCC2b by the WNK-SPAK signaling pathway.

Taken together, these data suggest that the N-terminal heterogeneity of KCC2 isoforms provides a mechanism for distinct regulation and developmental expression patterns. However, it remains unknown whether KCC2a maintains a key role in KCC2 function in the adult brain despite its low expression levels. If so, this isoform could be a prime target for new KCC2-modulating therapeutic compounds. Here, we investigated the age-dependent function of KCC2a in hippocampal and neocortical neurons. In primary hippocampal cultures, we found that endogenous KCC2a is enriched within dendritic KCC2 clusters, likely promoting its interaction with KCC2b. Knockdown of KCC2a in mature neurons revealed that KCC2a stabilizes KCC2b protein levels and expression at the cell membrane by preventing endocytosis and proteasomal degradation. Using KCC2a knockout mice, we demonstrated that this stabilizing function only prevails in mature neurons with low WNK-SPAK activity. In contrast, in immature neurons with high WNK-SPAK activity, KCC2a acts oppositely by promoting the phosphorylation of KCC2b at Thr1007, thereby reducing its membrane expression. This effect is associated with increased WNK1-mediated SPAK phosphorylation at Ser373. This suggests that KCC2a may recruit a functional protein unit, which includes both KCC2b and SPAK, to regulate chloride transport in dendrites. Taken together, our results reveal a novel mechanism by which KCC2a acts as a context-dependent regulator of KCC2b. Despite its weak expression in mature neurons, KCC2a transitions from a negative regulator to a stabilizing partner of KCC2b in response to developmental shifts in WNK-SPAK signaling.

## Results

### KCC2 isoform expression and subcellular localization in mature cortical neurons

First, we characterized and compared the expression of the two KCC2 isoforms, KCC2a and KCC2b, in maturing rat hippocampal neurons *in vitro* and in *ex vivo* samples from rat hippocampus and cortex. In both *in vitro* (from 7 to 24 DIV) and *ex vivo* (hippocampus and cortex from E18.5 to adult) samples, KCC2a mRNA levels remained stable throughout maturation (**Fig.1A-C**). In contrast, KCC2b expression increased significantly by ≍12-fold in the hippocampus and by ≍1.5-fold *in vitro* between 7 and 24 DIV (**Fig. 1B-C**). This differential regulation resulted in a marked decrease in the KCC2a/KCC2b mRNA ratio both *ex vivo* and *in vitro* (**Fig. 1B-C**). These results confirm that primary hippocampal cultures recapitulate the dynamic developmental shift in KCC2 expression, with KCC2a accounting for less than 5% of total KCC2 mRNA expression in mature neurons.

**Figure 1.**
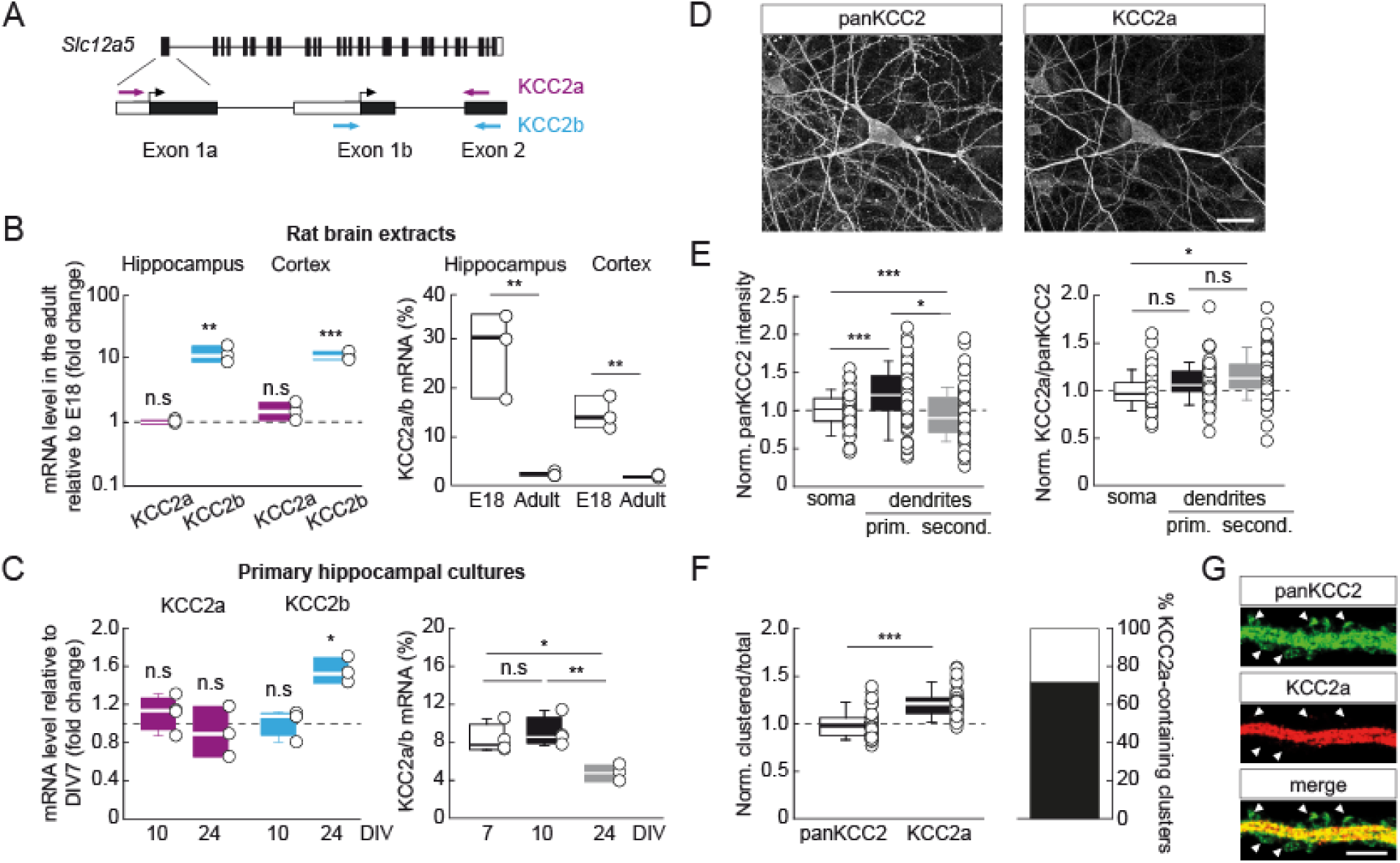
Expression pattern of KCC2 isoforms in rat hippocampal and cortical neurons. **A**, Schematic representation of the complementary sequences of KCC2a- and KCC2b-specific primers. **B**, Summary boxplots showing the quantitative real-time PCR analysis of KCC2a and KCC2b mRNA expression in the hippocampus and cortex of E18 and adult rats. Left: Isoform-specific mRNA expression relative to Ywhaz in adults, normalized to expression at E18. Right: KCC2a/KCC2b mRNA ratio in E18 embryos and in adults. n=3 animals/age and 3 technical replicates. Two-tailed t-test **p < 0.01; ***p < 0.001. **C**, Same as (B) in primary hippocampal cultures. Left: Isoform-specific mRNA expression relative to Ywhaz, normalized by expression at DIV7, one-way Kruskal-Wallis rank sum test and Dunn’s post-hoc test. Right: KCC2a/KCC2b mRNA ratio as a function of time *in vitro,* one-way ANOVA and Tukey post-hoc test. n=3-4 cultures and 3 technical replicates. *p < 0.05; **p < 0.01. **D**, Representative confocal maximum projection images of hippocampal pyramidal neurons immunostained with panKCC2 and KCC2a-specific antibodies. Scale: 25 µm. **E**, Summary boxplots showing the intensity of panKCC2 immunofluorescence (left, one-way Kruskal-Wallis rank sum test and Dunn’s post-hoc test) and the ratio of KCC2a to panKCC2 immunofluorescence (right, one-way ANOVA and Tukey post-hoc test), along primary (prim.) and secondary (second.) dendrites, normalized to the soma. n = 58 neurons from 2 cultures. *p < 0.05; ***p < 0.001. **F**, Left: summary boxplots showing the ratio of immunofluorescence intensity within clusters to total intensity in dendritic sections, for panKCC2 or KCC2a immunostainings, normalized to that of panKCC2. n = 46 neurons from 2 cultures, Mann-Whitney rank-sum test. ***p < 0.001. Right: bar plot showing the percentage of panKCC2 clusters colocalizing (black) or not (white) with KCC2a- immunostained clusters. **G**, Representative confocal images of a dendritic section immunostained with panKCC2 (green) and KCC2a (red) antibodies. Note the complete and specific lack of KCC2a immunostaining in dendritic spines (arrowheads). Scale: 5 µm.

Previous work in adult mouse brain suggested KCC2a may be preferentially enriched in distal dendrites compared to KCC2b (Markkanen *et al*, 2014). We therefore compared the subcellular localization of KCC2a and KCC2b in mature (24 DIV) hippocampal neurons using a panKCC2 antibody (recognizing both isoforms) and a KCC2a-specific antibody (Uvarov *et al*, 2009) (**Fig. S1**). PanKCC2 immunostaining was 20% stronger in primary dendrites compared to the soma (p<0.001; **Fig. 1E**). In addition, the KCC2a/panKCC2 fluorescence ratio was significantly increased in secondary dendrites compared to the soma (p<0.05; **Fig. 1E**), demonstrating that KCC2a is relatively enriched in dendrites compared to KCC2b. As previously demonstrated (Al Awabdh *et al*, 2022; Chamma *et al*, 2012; Gauvain *et al*, 2011), KCC2 immunostaining exhibited heterogeneity, with clusters present in both the soma and dendrites of hippocampal neurons. Notably, a greater fraction of KCC2a immunofluorescence was found within clusters compared to panKCC2 (p<0.001; **Fig. 1F**). Colocalization analysis showed that more than 70% of KCC2 clusters contain the KCC2a isoform (**Fig. 1F**). However, KCC2a-immunoreactivity was notably absent from dendritic spines (**Fig. 1G**), where KCC2 has been shown to form large clusters implicated in non-canonical, ion transport-independent functions (Gauvain *et al*, 2011). These results are consistent with an enrichment of KCC2a within subcellular domains critical for Cl^−^ extrusion and GABA signaling in mature hippocampal neurons (Raimondo *et al*, 2012).

### KCC2a controls KCC2b stability and function in mature hippocampal neurons

Despite its low expression levels, our data suggest that KCC2a may be optimally positioned to regulate chloride transport in mature hippocampal neurons. To evaluate the localization of each KCC2 isoform at the plasma membrane, we expressed recombinant KCC2a and KCC2b with a 3xFlag tag in their second extracellular loop, thereafter referred to as KCC2-Flag (Chamma *et al*, 2012; **Fig. 2A–B**). This enabled us to specifically detect KCC2 membrane pools using live immunostaining (**Fig. 2A-C**). The recombinant proteins showed no significant difference in total expression (p=0.206; **Fig. 2D**). However, the relative membrane expression of recombinant KCC2a was slightly but significantly lower than that of KCC2b (p<0.01; **Fig. 2D**), consistent with prior studies (Markkanen *et al*, 2014). These results suggest that, despite a small difference in membrane enrichment the overexpressed isoforms have similar inherent stability.

**Figure 2.**
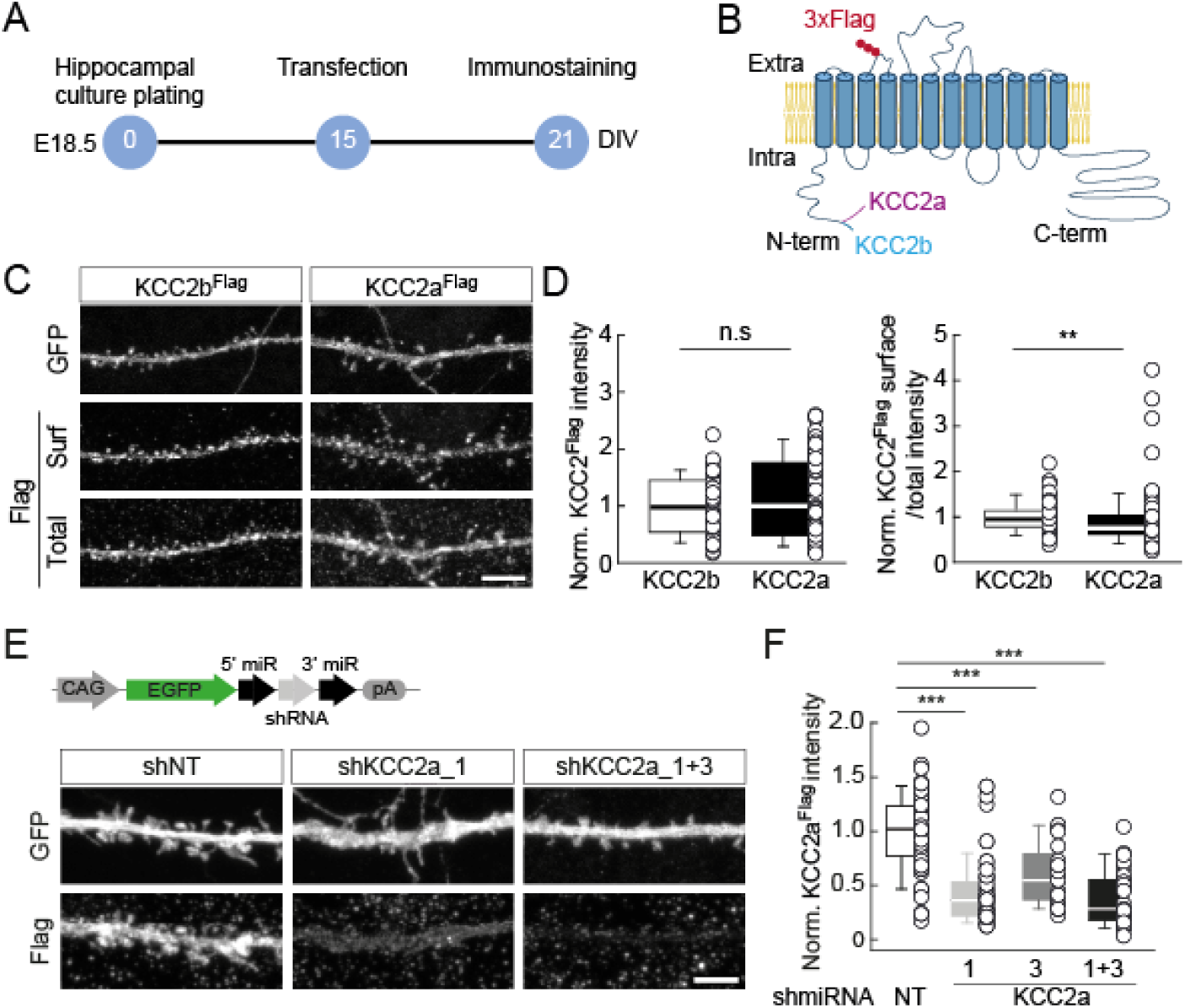
KCC2a-specific knockdown by RNA interference in mature hippocampal neurons. **A**, Experiment timeline. **B**, Schematic structure of the recombinant KCC2a-Flag and KCC2b-Flag proteins. **C**, Representative confocal maximum projection images of dendritic sections of hippocampal neurons transfected with plasmids encoding for GFP and either KCC2b-Flag or KCC2a-Flag. Neurons were immunostained for plasmalemmal and total Flag (see Methods) as well as GFP. Scale: 5 µm. **D**, Summary boxplots showing the immunofluorescence intensity of recombinant KCC2-Flag isoforms (left) and the ratio of membrane-inserted (surf.) to the total (total) KCC2-Flag immunofluorescence intensity normalized to that of KCC2b-Flag (right). KCC2b, n = 58; KCC2a, n = 65 cells from 5 independent cultures. Mann-Whitney rank-sum tests. **p < 0.01. **E**, Top: Design of the plasmids encoding the shRNAs targeting KCC2a. Bottom: Representative confocal maximum projection images of dendritic sections of neurons transfected with plasmids encoding either for a control scrambled shRNA (shNT) or KCC2a-specific shRNAs (shKCC2a_1 and shKCC2a_3), as well as a plasmid encoding recombinant KCC2a-Flag. Neurons were immunostained for Flag and GFP. Scale: 5 µm. **F**, Summary boxplot showing the immunofluorescence intensity of recombinant KCC2a-Flag, normalized to shNT. shNT, n = 68; shKCC2a_1, n = 65; shKCC2a_3, n = 25; shKCC2a1+3, n = 27 cells; from 5 (shNT and shKCC2a_1) or 2 (shKCC2a_3 and shKCC2a_1+3) independent cultures. One-way Kruskal-Wallis rank sum test and Dunn’s post-hoc test. ***p < 0.001.

To investigate the functional role of KCC2a in hippocampal neurons, we performed an isoform-specific knockdown using a combination of two shRNAs (shKCC2a_1+3). In HEK-293 cells expressing either recombinant KCC2a or KCC2b, these shRNAs specifically reduced KCC2a expression within 48 hours, with no significant effect on KCC2b (**Fig. S2**). In hippocampal neurons, chronic KCC2a knockdown (from DIV15) reduced recombinant KCC2a expression by over 60% (p<0.001; **Fig. 2E-F**). Surprisingly, despite KCC2a accounting for approximately 5% of total KCC2 mRNA, its suppression resulted in a 28% decrease in the total level of endogenous KCC2 protein (p < 0.001; **Fig. 3A-C**). This effect was accompanied by a significant decrease in the density of KCC2 clusters, as well as in their mean area and intensity (p ≤ 0.002 for all; **Fig. 3D**).

**Figure 3.**
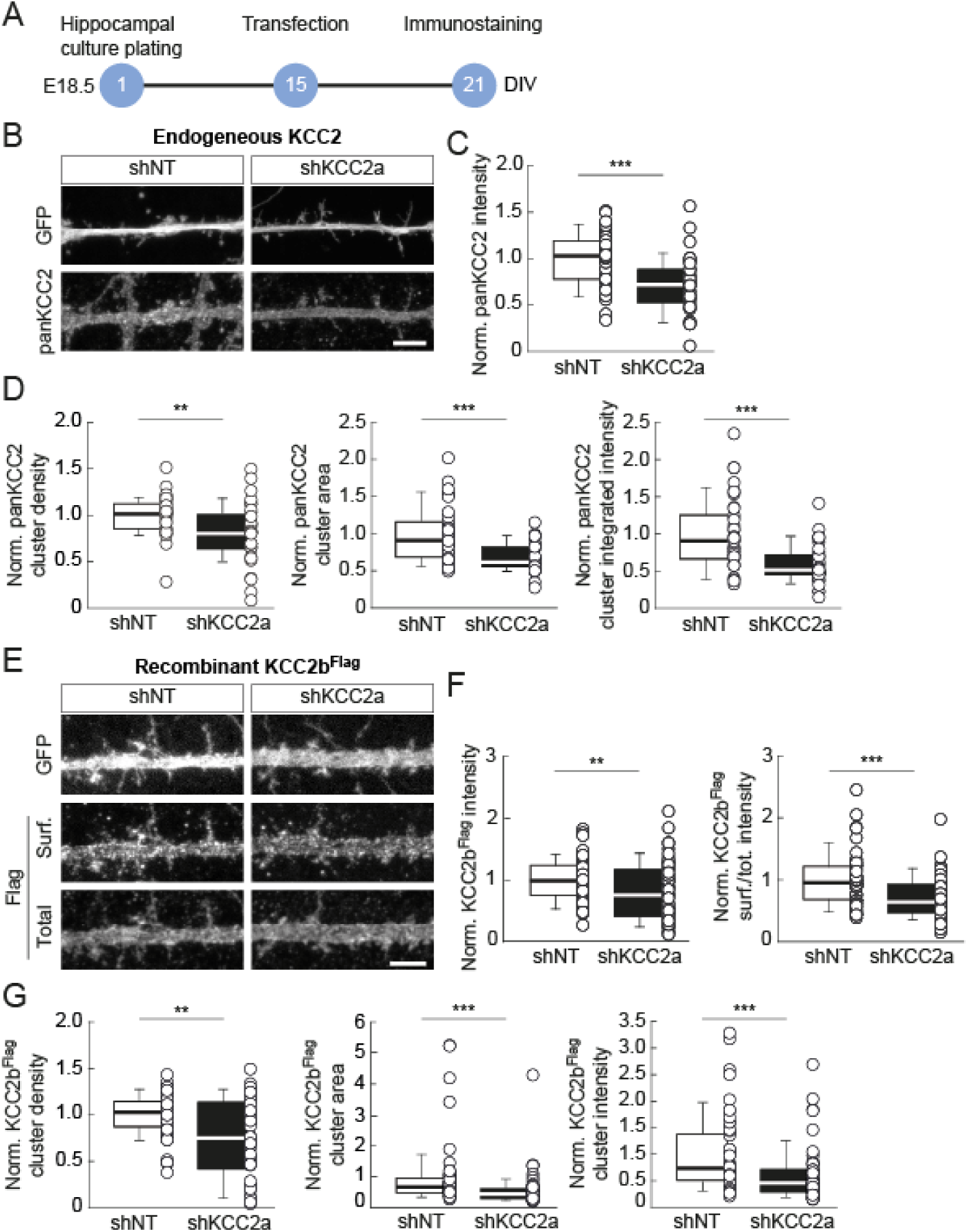
Specific KCC2a knockdown reduces KCC2b protein expression, membrane expression, and clustering. **A**, Experiment timeline. **B**, Representative confocal maximum projection images of dendritic sections of neurons transfected with plasmids encoding scrambled (shNT) or KCC2a-specific shRNAs (shKCC2a), immunostained for endogenous KCC2 and GFP. Scale: 5 µm. **C**, Boxplot showing the intensity of endogenous KCC2 immunofluorescence in neurons transfected with shNT or shKCC2a_1+3. Two-tailed t-test. ***p < 0.001. **D**, Boxplots showing the properties of endogenous panKCC2-immunopositive clusters. Left: cluster density/10µm^2^, middle: mean cluster surface, right: mean integrated intensity of cluster fluorescence. shNT, n = 35; shKCC2a, n = 35 cells, from 3 independent cultures. Mann-Whitney rank-sum tests. **p < 0.01; ***p < 0.001. **E**, Same as (B) with neurons expressing recombinant KCC2b-Flag and immunostained for Flag and GFP. Scale: 5 µm. **F**, Boxplots showing the intensity of recombinant KCC2b-Flag immunofluorescence (left) and the ratio of plasmalemmal (surf.) to the overall (total) Flag immunofluorescence intensity (right). Mann-Whitney rank-sum tests. **p < 0.01; ***p < 0.001. **G**, Same as (D), for neurons co-transfected with recombinant KCC2b-Flag and shRNA-expressing plasmids. shNT, n = 53; shKCC2a, n = 54 cells; from 4 independent cultures. Mann-Whitney rank-sum tests. **p < 0.01, ***p < 0.001. All data are normalized to shNT.

These results suggested that KCC2a controls the expression of KCC2b in mature hippocampal neurons. To test this hypothesis, we measured recombinant KCC2b expression upon KCC2a knockdown (**Fig. 3E-G**). Consistent with its effects on endogenous KCC2, KCC2a knockdown reduced the expression of recombinant KCC2b as well as its relative membrane enrichment by 20% and 30%, respectively (p≤0.01; **Fig. 3F**). This reduction was accompanied by a significant decrease in KCC2b cluster density, area, and fluorescence intensity (p≤0.003 for all; **Fig. 3G**). Together, these data demonstrate that KCC2a exerts strong, non-stoichiometric control over KCC2b protein expression, membrane availability, and clustering.

Since membrane stability and clustering are essential for KCC2 ion transport function (Chamma *et al*, 2012; Chamma *et al*, 2013; Côme *et al*, 2019), we investigated whether KCC2a regulates transmembrane chloride transport in hippocampal neurons. We assessed neuronal chloride extrusion efficiency by measuring the difference in the GABA_A_R reversal potential (E_GABA_) between the soma and the dendrite (ΔE_GABA_) (Al Awabdh *et al*, 2022; Gauvain *et al*, 2011; Khirug *et al*, 2008) (**Fig. 4A-D**). As expected, KCC2a knockdown did not significantly impact somatic E_GABA_ (p=0.8; **Fig. 4E**). However, it resulted in a 41% decrease in ΔE_GABA_ (6.0±1.1 versus 3.2±0.7 mV/100μm; p<0.05; **Fig. 4F**), indicating a significant reduction in KCC2-mediated chloride export along dendrites. Taken together, our results demonstrate the unsuspected role of the minor KCC2a isoform in controlling both KCC2b membrane expression and function in mature hippocampal neurons.

**Figure 4.**
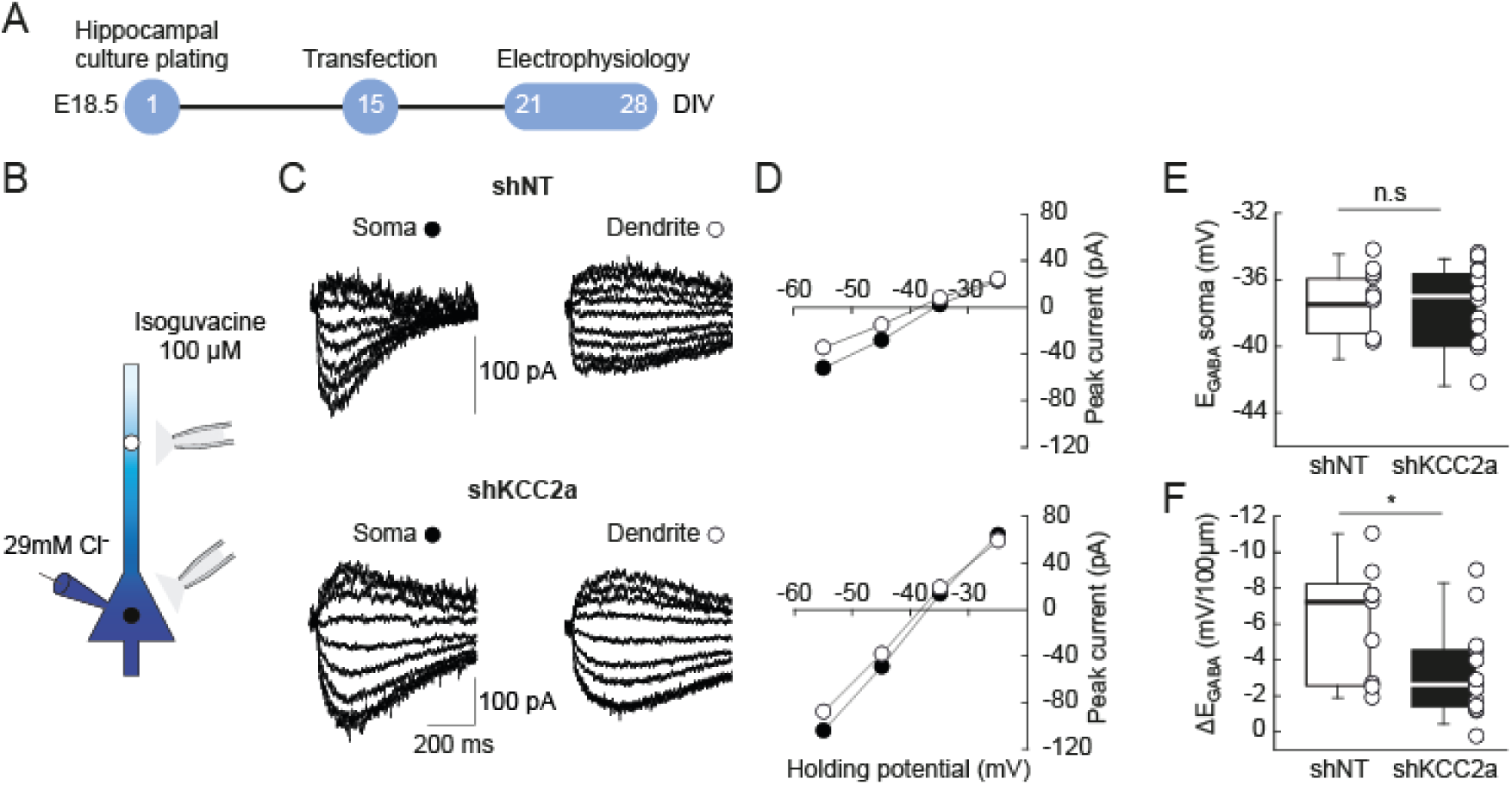
Specific KCC2a knockdown impairs chloride export in hippocampal neurons. **A**, Experiment timeline. **B**, Schematic representation of the experiment. Isoguvacine (100 µM) was locally released onto the soma or dendrite of neuron in whole-cell patch-clamp configuration. The patch pipette was filled with a high-chloride (29 mM) intracellular solution. **C**, Representative recordings of isoguvacine-evoked currents at varying potentials. Currents were evoked in neurons transfected with plasmids encoding either scrambled (shNT) or KCC2a-specific (shKCC2a) shRNAs**. D**, I–V curves corresponding to traces shown in C. **E, F,** Summary boxplots showing the reversal potential of isoguvacine-evoked currents at the soma (E_GABA_ soma, E) and the somato-dendritic E_GABA_ gradient (ΔE_GABA_, F) in transfected neurons expressing either shNT or shKCC2a_1+3. shNT, n = 9; shKCC2a, n = 14 cells; from 5 independent cultures. Two-tailed t-tests. *p < 0.05.

### KCC2a stabilizes KCC2b by hindering clathrin-mediated endocytosis and proteasomal degradation

Next, we explored the molecular mechanisms by which KCC2a stabilizes KCC2b in hippocampal neurons. Previous studies have shown that KCC2b undergoes constitutive internalization via a dynamin- and clathrin-dependent pathway (Chamma *et al*, 2013; Friedel *et al*, 2017; Zhao *et al*, 2008). Therefore, we investigated the involvement of KCC2a in KCC2b endocytosis. To this end, we blocked clathrin-dependent endocytosis using a dynamin inhibitory peptide (DiP) (Kittler *et al*, 2000) for 40 minutes prior to and during the 20-minute live staining (**Fig. 5A-C**). In neurons expressing a scrambled shRNA, dynamin inhibition resulted in a modest, though not significant, increase in recombinant KCC2b membrane expression (p=0.26; **Fig. 5C**). However, KCC2a knockdown significantly decreased the relative membrane expression of recombinant KCC2b (p<0.05; **Fig. 5C**). This reduction was fully reversed in KCC2a knocked-down neurons treated with DiP (p=0.26 compared to shNT+DiP; **Fig. 5C**). These results demonstrate that KCC2a stabilizes KCC2b at the membrane by preventing its clathrin-mediated endocytosis.

**Figure 5.**
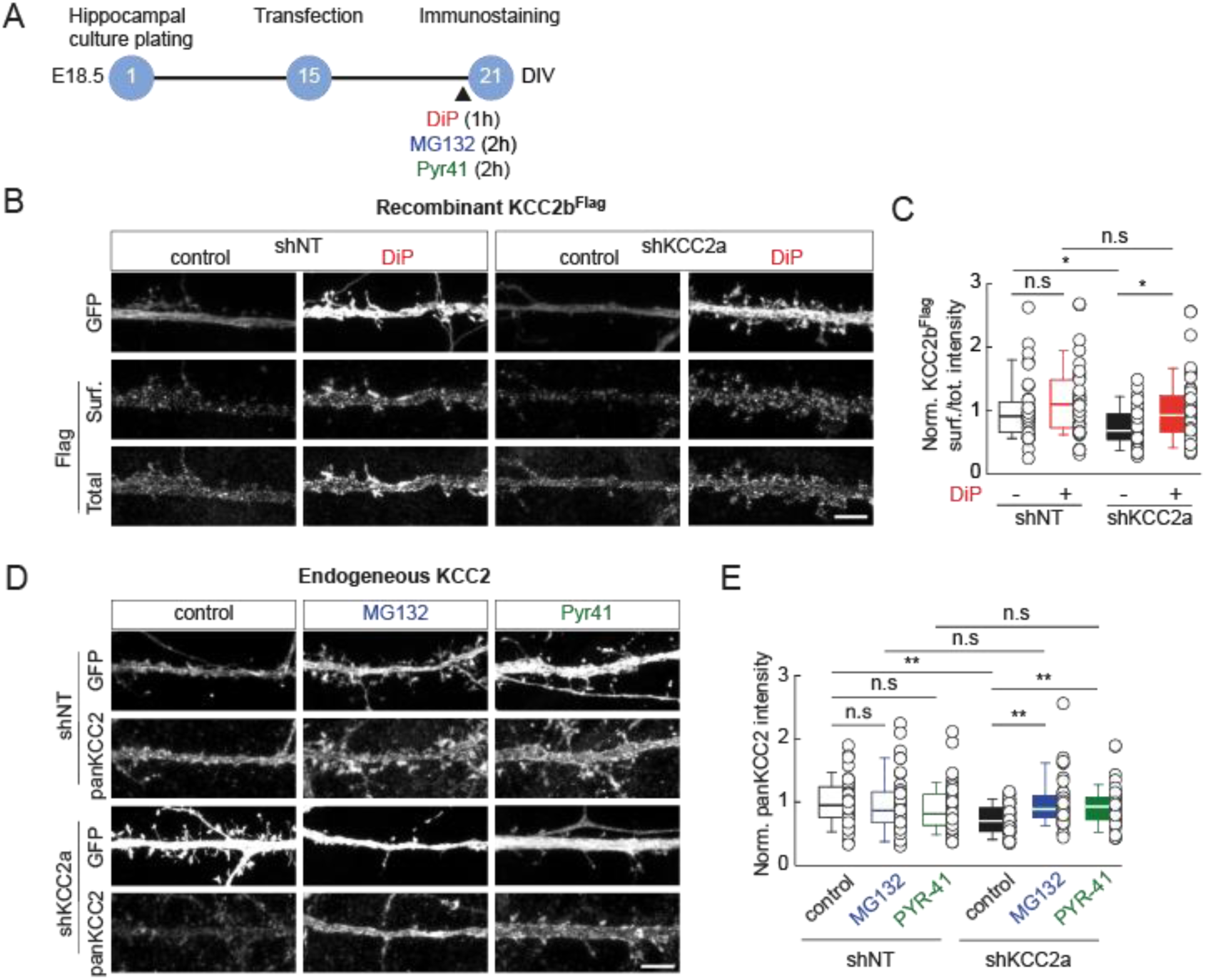
KCC2a prevents KCC2b endocytosis and degradation via the proteasomal pathway. **A**, Timeline of the experiments. **B**, Representative confocal maximum projection images of dendritic sections of mature hippocampal neurons expressing scrambled (shNT) or KCC2a-specific (shKCC2a) shRNA, treated with a dynamin inhibitory peptide (DiP, 50 µM) 40 minutes before and during live Flag immunostaining. Scale: 5 µm. **C**, Summary boxplot showing the ratio of plasmalemmal (surf.) to the overall (total) KCC2b-Flag immunofluorescence intensity, normalized to control shNT condition. shNT control, n = 37; shNT +DiP, n = 39; shKCC2a control, n = 42; shKCC2a +DiP, n = 41 cells, from 4 independent cultures. Mann-Whitney rank-sum tests, p-value corrected for multiple comparisons using the Holm-Šídák method. *p < 0.05. **D**, Representative confocal maximum projection images of dendritic sections of hippocampal neurons expressing scrambled (shNT) or KCC2a-specific shRNA. Two hours before immunostaining with a panKCC2 antibody, neurons were treated with either a proteasome blocker (MG132, 2.5 µM), an inhibitor of the ubiquitin-activating enzyme (PYR-41, 50 µM), or untreated (control). Scale: 5 µm. **E**, Summary boxplot showing the intensity of endogenous KCC2 immunofluorescence in the different conditions, normalized to shNT control. shNT control, n = 44; shNT +MG132, n = 38; shNT +PYR41, n = 43; shKCC2a control, n = 43; shKCC2a +MG132, n = 40; shKCC2a +PYR41, n = 44 cells, from 4 independent cultures. Whitney rank-sum tests, p-value corrected for multiple comparisons using the Holm-Šídák method. **p < 0.01.

KCC2 may be subject to recycling at the plasma membrane following endocytosis (Côme *et al*, 2019). Alternatively, it may undergo degradation through the lysosomal (Lee *et al*, 2010) or ubiquitin-proteasomal pathway (Hu *et al*, 2023). Next, we investigated whether KCC2a may be preventing KCC2 degradation, which could account for its reduced total expression upon KCC2a knockdown. Under basal conditions (scrambled shRNA), specifically blocking the proteasome with MG132 (2.5 µM) or polyubiquitination with PYR-41 (50 µM) for two hours did not significantly reduce endogenous KCC2 expression (p=0.85 and 0.41 respectively) (**Fig. 5D-E**). However, the decrease in endogenous KCC2 expression after KCC2a knockdown was fully abolished when either the proteasome or polyubiquitination was blocked (p>0.85 for both, compared to shNT with the corresponding drug; **Fig. 5E**). These results suggest that KCC2a controls KCC2 protein levels by preventing its polyubiquitination and subsequent degradation by the proteasome.

### KCC2a couples KCC2b to the WNK-SPAK regulatory pathway

The expression and function of the plasmalemmal KCC2 pool are tightly regulated by various mechanisms, including protein-protein interactions, calpain-mediated cleavage, and the phosphorylation of key residues in its C-terminal domain (Côme *et al*, 2019; Hartmann & Nothwang, 2022; Kahle *et al*, 2013; Pressey *et al*, 2022). In particular, the WNK-SPAK kinase pathway modulates KCC2 membrane stability in a chloride-dependent manner through phosphorylation of its Thr1007 residue (de Los Heros *et al*, 2014; Friedel *et al*, 2015; Heubl *et al*, 2017). KCC2a, but not KCC2b, was shown to contain a conserved binding motif for SPAK and co-immunoprecipitate with SPAK in heterologous cells (Uvarov *et al*, 2009; Markkanen *et al*, 2017). We therefore asked whether KCC2a is required for the WNK-SPAK-dependent modulation of KCC2b.

First, we examined protein extracts from the brains (excluding the brainstem) of KCC2a-knockout (KO) mice at postnatal day 9 (P9) (Markkanen *et al*, 2014), a developmental stage at which high WNK1-SPAK activity has been reported (Friedel *et al*, 2015). While the total level of KCC2 protein expression did not differ significantly from that of wild-type (WT) littermates (p=0.2; **Fig. 6A-B**), KCC2a-KO mice exhibited a 62% decrease in KCC2 Thr1007 phosphorylation (p<0.001; **Fig. 6A-B**). This suggests that KCC2a may be necessary for KCC2 phosphorylation by SPAK. Interestingly, KCC2a-KO mice also showed a 43% decrease in SPAK Ser373 phosphorylation (p<0.05; **Fig. 6C-D**), but no changes in WNK1 Ser382 phosphorylation (**Fig. S3**). This suggests that KCC2a may promote WNK1-dependent SPAK phosphorylation in the immature brain without affecting WNK1 activity itself. By contrast, no changes in KCC2 or SPAK expression or phosphorylation were observed in the brains of adult KCC2a-KO mice, in which basal WNK1-SPAK pathway activity is low (Friedel *et al*, 2017; Heubl *et al*, 2017) (p>0.2; **Fig. 6E-H**). Together, these data show that KCC2a regulates SPAK activity and KCC2b Thr1007 phosphorylation only in the immature brain.

**Figure 6.**
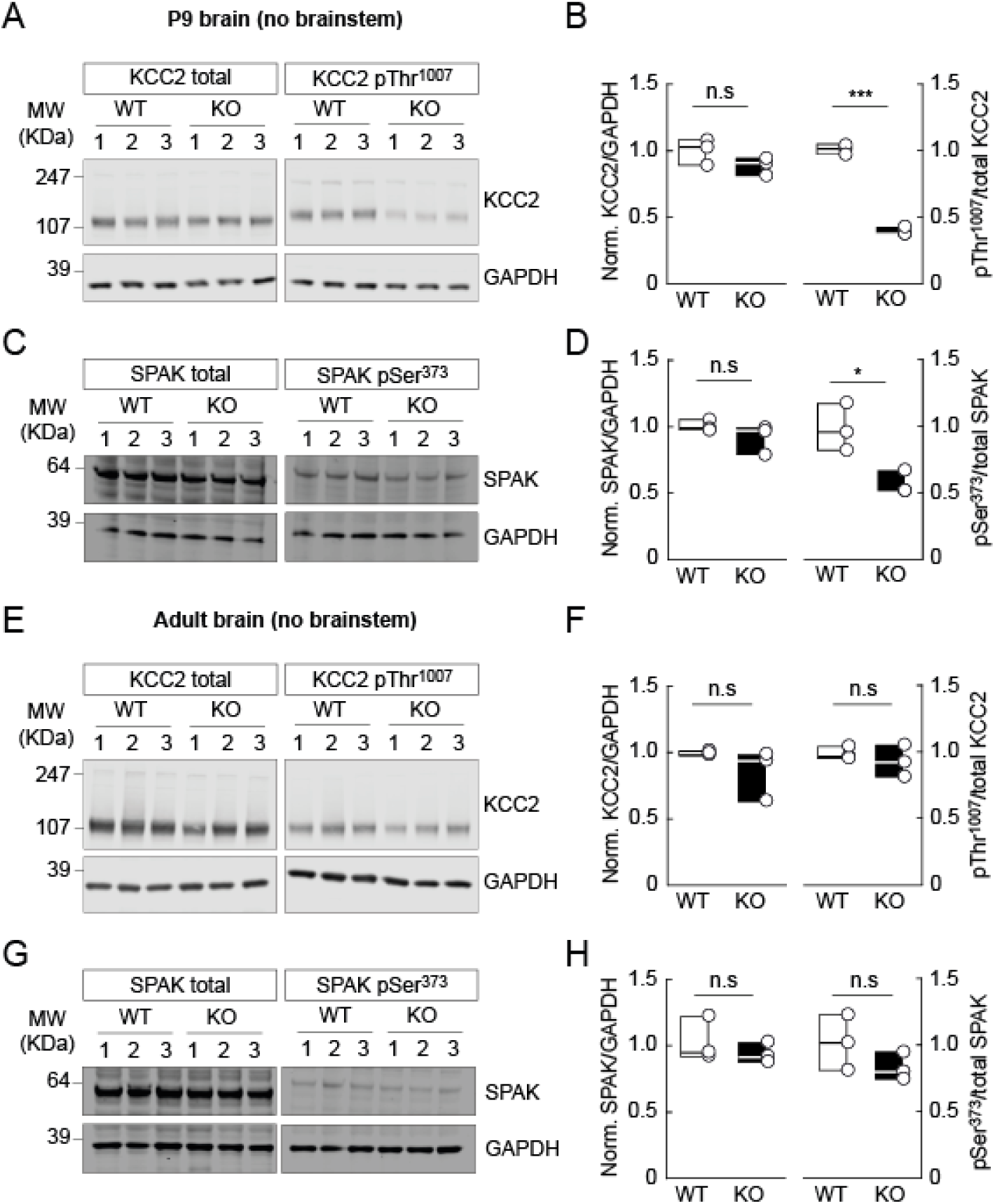
KCC2a gates WNK/SPAK-mediated KCC2 Thr1007 phosphorylation in the immature, but not adult postnatal brain. **A**, **C,** Western blots of brain extracts (without brainstem) from P9 KCC2a-KO pups and WT littermates. Each membrane was probed for GAPDH, as well as either endogenous KCC2, KCC2 Thr^1007P^, SPAK or SPAK Ser^373P^. **B**, **D**, Summary boxplots showing the ratio of total panKCC2 or SPAK immunofluorescence to that of GAPDH (left) or phosphorylated KCC2 or SPAK immunofluorescence to that of GAPDH and normalized by the corresponding total protein (right). All results are normalized to WT. WT, n = 3; KO, n = 3 animals; 3 technical replicates from the same brain extracts. Two-tailed t-test. *p < 0.05; ***p < 0.001. **E, G**, Same as (A, C) for adult KCC2a-KO and WT littermates. **F, H**, Same as (B, D) for adult brain extracts. WT, n = 3; KO, n = 3 animals; 3 technical replicates from the same brain extracts. Two-tailed t-test.

Next, we investigated whether this age-dependent effect was due to different levels of KCC2a expression or WNK-SPAK activity in immature versus adult brains. To explore this possibility, we examined the impact of KCC2a knockdown on WNK-SPAK-dependent regulation of KCC2b in mature neurons under chronic activation of the WNK-SPAK pathway. We used a TetOn system, which allows for precise temporal control, to overexpress a constitutively active WNK1 bearing the S382E mutation (WNK1-CA) (Friedel *et al*, 2015; Inoue *et al*, 2012) for 24 hours in mature hippocampal neurons in culture (**Fig. 7A-B**). In line with prior studies (Friedel *et al*, 2015; Heubl *et al*, 2017), we observed a 30% decrease in recombinant KCC2b membrane expression in mature neurons with intact KCC2a expression when WNK1-CA expression was induced by doxycycline (p<0.05; **Fig. 7D**). In contrast, in KCC2a-knockdown neurons, the WNK1-CA-induced decrease in KCC2b-Flag membrane expression was abolished and even reversed, leading to a 55% increase in membrane expression (p<0.01; **Fig. 7F**). This demonstrates that SPAK-dependent phosphorylation of KCC2b is tightly dependent on KCC2a, and that this effect is driven by WNK1 pathway activity rather than by the relative expression level of KCC2a.

**Figure 7.**
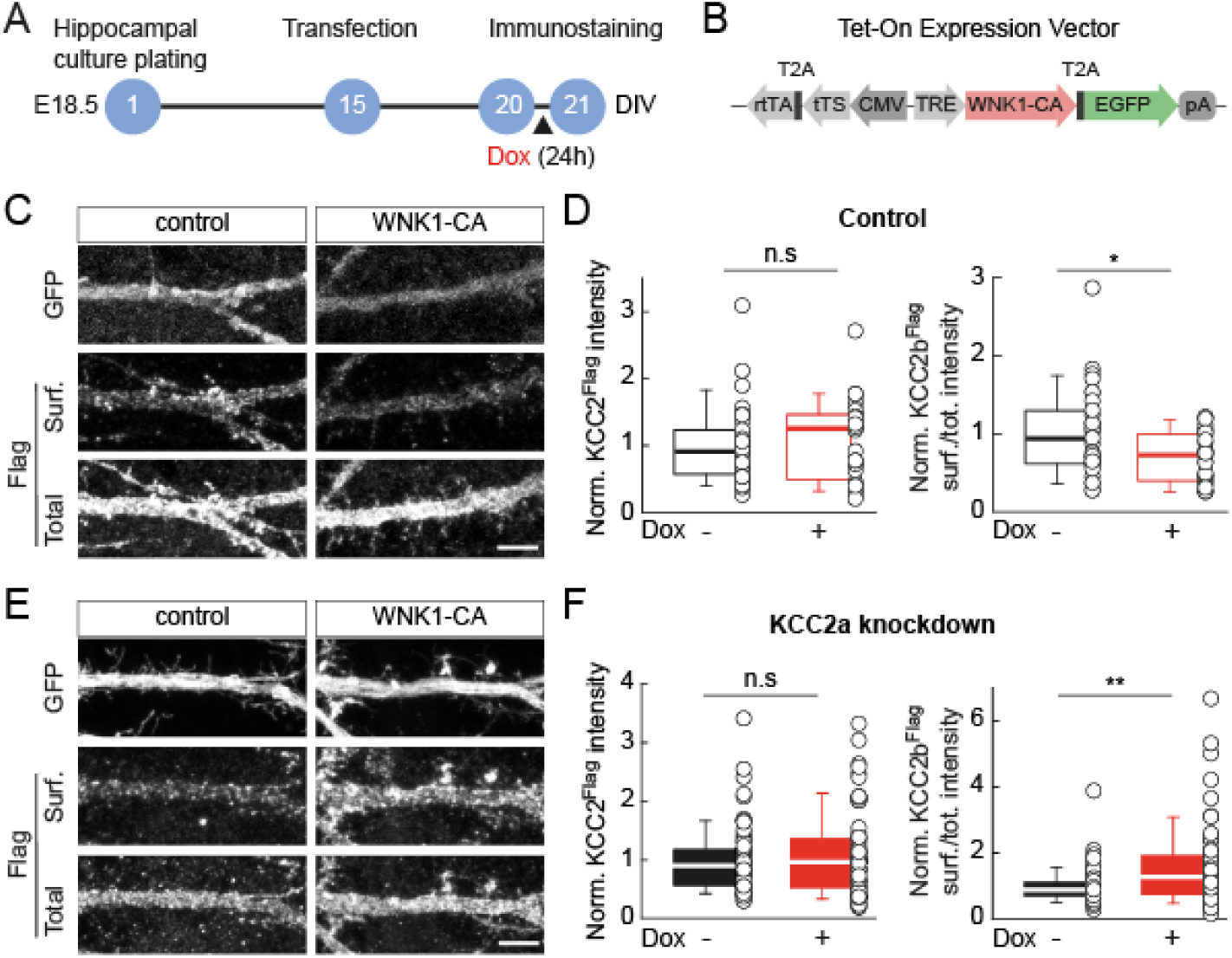
KCC2a gates WNK1-dependent regulation of KCC2b membrane expression in mature hippocampal neuron. **A**, Experiment timeline. **B**, Design of the plasmid encoding for the constitutively activated WNK1 (WNK1-CA), bearing a S382E mutation. **C**, Representative confocal maximum projection images of dendritic sections of hippocampal neurons transfected with plasmids encoding GFP, WNK1-CA and recombinant KCC2b-Flag. Doxycycline (Dox+, 2 µg/ml) was added to culture medium 24h before live immunostaining. Neurons were immunostained for Flag and GFP. Scale: 5 µm. **D**, Summary boxplots showing the intensity of Flag immunofluorescence (left) and the ratio of plasmalemmal (surf) to the overall (total) Flag immunofluorescence intensity (right) in neurons treated or not with doxycycline. Dox-, n = 31; Dox+, n = 28 cells; from 3 independent cultures. Mann-Whitney rank-sum tests. *p < 0.05. **E**, Same as (C), in neurons additionally co-transfected with plasmids encoding shKCC2a-specific shRNAs. Scale: 5 µm. **F**, Same as (D), in KCC2a-specific knocked-down neurons. Dox-, n = 71; Dox+, n = 67 cells; from 6 independent cultures. Mann-Whitney rank-sum tests. **p < 0.01. All data were normalized to control (no doxycycline).

## Discussion

In this study, we reveal the critical and previously unsuspected role of the minor KCC2a isoform in regulating the expression, membrane stability, and function of the predominant KCC2b isoform in mature hippocampal neurons. Despite accounting for less than 5% of total KCC2 mRNA in these neurons, we demonstrate that KCC2a preferentially accumulates in dendritic KCC2 clusters and exerts powerful, age-dependent, and non-stoichiometric control over KCC2b. Suppressing KCC2a in mature hippocampal neurons significantly reduces total KCC2 protein levels and impairs dendritic chloride extrusion. Mechanistically, we demonstrate that KCC2a promotes KCC2b clustering and prevents its clathrin-mediated endocytosis and subsequent proteasomal degradation. Furthermore, we establish that KCC2a provides an essential link between KCC2 and the WNK-SPAK signaling pathway, promoting KCC2b Thr1007 phosphorylation when this cascade is active. Therefore, in immature neurons, KCC2a contributes to maintaining low KCC2b membrane expression. Together, these findings revise the current model of KCC2 regulation. They identify the minor KCC2a isoform as a master regulator that controls the stability and function of KCC2 as well as the formation of a WNK1-SPAK-KCC2 complex that gates WNK1-mediated control of KCC2 phosphorylation.

Consistent with previous studies (Markkanen *et al*, 2017; Uvarov *et al*, 2007), our results indicate that KCC2a accounts for a small fraction (<10%) of total KCC2 transcripts in mature hippocampal neurons *in vivo* and in primary cultures. Although it is difficult to extrapolate transcript relative abundance to protein levels, western blot data support that KCC2a may represent a similar proportion of total KCC2 in the cortex and hippocampus (Uvarov *et al*, 2009). Therefore, our observations raise the question of how such a weakly expressed isoform may significantly impact the membrane stability and function of the total KCC2 pool. Our data suggest that this may, at least in part, reflect the preferential targeting of KCC2a to functional KCC2 clusters. While KCC2 is expressed throughout the somato-dendritic membrane and exhibits both a diffuse and a clustered distribution, several studies have shown that KCC2 clusters likely represent the most transport-active form of the protein (Al Awabdh *et al*, 2022; Chamma *et al*, 2013; Côme *et al*, 2019; Gauvain *et al*, 2011; Watanabe *et al*, 2009). Indeed, manipulations of KCC2 clustering result in changes in chloride export function, even with no change in the total plasmalemmal fraction (Al Awabdh *et al*, 2022; Watanabe *et al*, 2009). Despite a slightly lower plasmalemmal-to-total ratio of recombinant KCC2a compared to KCC2b (Fig. 2D), which is consistent with previous work (Markkanen *et al*, 2014), our data show that endogenous KCC2a is preferentially expressed in proximal and distal dendrites compared to KCC2b (Fig. 1E). Additionally, KCC2a displays an increased cluster-to-total ratio, and the vast majority (>70%) of KCC2 clusters contain KCC2a (Fig. 1F). Thus, despite its low relative mRNA expression level, KCC2a subcellular expression pattern may be optimal for influencing KCC2b clustering and function. Previous studies have highlighted the presence of large KCC2 clusters within dendritic spines (Chamma *et al*, 2013; Gauvain *et al*, 2011; Gulyas *et al*, 2001). There, KCC2 was shown to influence spine maturation and volume, actin dynamics, and synaptic function and plasticity (Awad *et al*, 2018; Chevy *et al*, 2015; Gauvain *et al*, 2011; Li *et al*, 2007; Llano *et al*, 2015). Remarkably, we now show that endogenous KCC2a-immunoreactivity is completely absent from dendritic spines (Fig. 1G). This suggests that the two isoforms may be differentially targeted to specific subcellular compartments and that only KCC2b may be endowed with non-canonical functions in dendritic spines.

Despite the imbalance in KCC2 isoform stoichiometry, our data reveal that KCC2a plays a critical and unexpected role in controlling KCC2 in mature hippocampal neurons. Our experiments on neurons expressing recombinant Flag-tagged KCC2b revealed trans-regulation of KCC2 isoforms: KCC2a knockdown reduced KCC2b total expression, plasmalemmal fraction, and clustering (Fig. 3), which was associated with reduced chloride export function (Fig. 4). While KCC2 was initially suggested to undergo very rapid turnover (Rivera *et al*, 2004), subsequent studies revealed that it is long-lived, with stability under physiological conditions ranging from hours to days (Puskarjov *et al*, 2012). However, KCC2 can be modulated by activity within minutes. This modulation involves the phosphorylation or dephosphorylation of key residues in its C-terminal domain or calpain cleavage. These changes result in increased membrane diffusion, endocytosis, and degradation by the lysosomal and proteasomal pathways (Côme *et al*., 2019). Our data show that, under basal conditions, blocking clathrin-dependent endocytosis for one hour or polyubiquitination and proteasomal degradation for two hours has no significant impact on the plasmalemmal fraction or total expression of KCC2, respectively (Fig. 5). These results are inconsistent with previous observations of rapid recycling of membrane-inserted KCC2 (Zhao *et al*, 2008; Lee *et al*, 2010), but they do confirm the relative stability of the total KCC2 pool within this time range. However, we demonstrate that these manipulations completely reverse the effects of chronic KCC2a knockdown. This suggests that KCC2a acts to promote KCC2 membrane stability and prevent its ubiquitination and proteasomal degradation, even under basal activity conditions. The molecular determinants of this stabilizing effect remain to be established, but they could involve KCC2 interactions with its numerous partner proteins (Al Awabdh *et al*, 2022; Mahadevan *et al*, 2017; Smalley *et al*, 2020). In particular, the proteasomal pathway has recently been identified as a key element in KCC2 degradation, involving the direct interaction of KCC2b with FBXL4, a protein that forms the E3 ubiquitin ligase complex SCF-FBXL4 and that is essential for KCC2 polyubiquitination (Hu *et al*, 2023; Chen *et al*, 2025). KCC2a may hinder this interaction, thereby preventing KCC2b ubiquitination and degradation. Alternatively, KCC2a may influence KCC2b interaction with partners known to control its membrane stability and/or clustering, such as gephyrin (Al Awabdh *et al*, 2022), SNARE proteins (Asraf *et al*, 2022), Neto2 (Ivakine *et al*, 2013), and CKB (Inoue *et al*, 2006).

In heterologous cells, only KCC2a, and not KCC2b, has been shown to interact with SPAK (Markkanen *et al*, 2017), an effector of WNK and a major regulator of cation-chloride cotransporters (Alessi *et al*, 2014). SPAK has also been shown to suppress chloride transport mediated by recombinant KCC2a, but not KCC2b (Markkanen *et al*, 2017). These observations are consistent with the presence of a SPAK-binding consensus motif in the N-terminal domain of KCC2a (Uvarov *et al*, 2007). Therefore, SPAK may interact exclusively with KCC2a, and/or KCC2a may be necessary for SPAK-dependent modulation of KCC2b. Indeed, previous work has revealed that KCC2a can oligomerize with KCC2b (Uvarov *et al*, 2009). Our *in vitro* and *in vivo* results now demonstrate that the KCC2a isoform is crucial for SPAK-dependent phosphorylation and internalization of KCC2b. Endogenous phosphorylation of the KCC2 Thr1007 residue was strongly reduced in brain extracts from P9 KCC2a-KO pups (Fig. 6B), a developmental stage at which WNK/SPAK activity is high (Friedel *et al*, 2015). Additionally, KCC2a knockdown suppressed the downregulation of recombinant KCC2b membrane expression induced by constitutive WNK1 activation in mature hippocampal neurons (Fig. 7). These results demonstrate that KCC2a is necessary for WNK-SPAK-dependent regulation of KCC2 and that this effect is not strictly age-dependent but rather context-specific and persists in mature neurons when the WNK-SPAK pathway is reactivated. More surprisingly, our data show that KCC2a influences WNK-dependent SPAK activation, as evidenced by reduced SPAK phosphorylation on its Ser373 residue, but not WNK1 phosphorylation, in KCC2a-KO brain extracts (Fig. 6 and Fig. S3). These findings suggest the possibility that interaction with KCC2a may somehow expose SPAK to WNK-mediated phosphorylation, thereby promoting its activation. Collectively, these data support a model in which KCC2a is essential for assembling a functional unit comprising KCC2b and the regulatory kinase SPAK, which controls chloride transport and its regulation in cortical neurons.

Given the versatile and fine-tuning role of KCC2a in regulating KCC2b, it is somewhat surprising that KCC2a KO mice were reported to display only a mild phenotype, with a low breathing rate and abnormal occurrence of apnea at birth (Dubois *et al*, 2018). Our experiments using chronic KCC2a knockdown *in vitro* predict a reduced KCC2 expression and function in the adult forebrain of KCC2a KO mice, which are expected to impact neuronal function and cortical rhythmogenesis (Goutierre *et al*, 2019; Simonnet *et al*, 2023). In support of a specific role for this isoform in pathology, KCC2a was also found to be specifically reduced in brain samples of patients with Rett syndrome (Hinz *et al*, 2019). However, compensatory mechanisms may be at play in the constitutive KO mice to mask these effects. Further phenotypic exploration of these mice may be needed to reveal anomalies related to altered KCC2 function. Our data, on the other hand, also predict that acute neuronal stress induced under conditions associated with re-activation of the WNK-SPAK pathway such as *status epilepticus*, stroke (Bhuiyan *et al*, 2022) or brain trauma (Gong *et al*, 2021), may differentially affect WT versus KCC2a KO mice. Since KCC2a appears to gate WNK-SPAK-dependent KCC2 downregulation, KO mice may display increased resilience or a better outcome in these conditions. Therefore, clinically, compounds that specifically prevent KCC2a-SPAK interaction may represent useful adjunct therapeutics for *status epilepticus*, stroke, and other conditions.

## Material and methods

### Animal experimentation

All animal procedures were carried out according to the European Community Council directive of 24 November 1986 (86/609/EEC) and the guidelines of the French Agriculture and Forestry Ministry for handling animals (decree 87-848) and were approved by the “Direction Départementale de la Protection des Populations de Paris” (Institut du Fer à Moulin animal facility, license E750522). Brains from KCC2a-KO mice and their WT littermates were kindly provided by Muriel Thoby-Brisson’s team and were handled as in (Dubois *et al*, 2018).

### Primary hippocampal cultures

Primary hippocampal neuron cultures were prepared from the hippocampi of Sprague Dawley (Rj:SD) rats on embryonic day 18.5 (E18.5), as previously described (Al Awabdh *et al*, 2022; Gauvain *et al*, 2011). The hippocampi were dissociated using 0.25% trypsin (Gibco) and then triturated mechanically in ice-cold HBSS (Gibco) supplemented with HEPES (Thermo Fisher Scientific). The neurons were plated at a density of 1.5×10⁵ cells per 18-mm glass coverslip pre-coated with 50 μg/ml poly-D,L-ornithine (Sigma-Aldrich). The initial plating medium consisted of minimum essential medium (Gibco) supplemented with 10% horse serum (Gibco), 2 mM L-glutamine (Thermo Fisher Scientific), and 1 mM sodium pyruvate (Thermo Fisher Scientific). After a three to four-hour attachment period, the plating medium was replaced with a serum-free culture medium composed of Neurobasal medium (Gibco) supplemented with B27 (Gibco), 2 mM L-glutamine (Thermo Fisher Scientific), and antibiotics (200 units/mL penicillin and 200 μg/mL streptomycin, Gibco). The neurons were maintained in an incubator at 37 °C with 5% CO₂ for up to four weeks, with one-third of the medium replaced weekly.

### HEK-293 cell cultures

HEK-293 cells were maintained in high glucose (4.5g) DMEM supplemented with GlutaMAX and pyruvate (Gibco), 10% FBS (Gibco), and 0.2% penicillin/streptomycin (Gibco). For transfection, cells were plated at a density of 5×10^4^ cells per 18-mm glass coverslip in 12-well plates without antibiotics.

### Plasmid constructs

The KCC2b-Flag construct was generated by inserting a sequence encoding a 3xFlag tag (DYKDHDGDYKDHHIDYKDDDD) into the second extracellular loop of KCC2, between the NsiI and NheI restriction sites of the full-length pEGFP–IRES–KCC2 construct (GenBank accession number U55816) (Chamma *et al*, 2013). The KCC2a-Flag construct was subcloned by inserting a synthetic 120-bp fragment encoding the KCC2a-specific N-terminus into the KCC2b-Flag backbone to replace the KCC2b-specific N-terminal coding sequence (Genscript, sequence available upon request). Specific knockdown of rat KCC2a was achieved using two distinct shRNAs (short hairpin RNAs) embedded in a miR-30 backbone obtained from VectorBuilder. The target shRNA sequences, which were designed using VectorBuilder and Thermo Fisher Scientific BLOCK-iT™ RNAi Designer tools, were: CTCCCGAGGGAAGACGTCAAA (shKCC2a_1), ACCCAGAGTCCCGCCGGCATTCGGT (shKCC2a_3). The scrambled shRNA sequence was: ACCTAAGGTTAAGTCGCCCTCG.

The WNK1-CA construct expresses a constitutively active form of rat WNK1 with an S382E mutation, which is cloned into a Tet-On backbone (TRE>WNK1:T2A:GFP-rev(CMV>tTS:t2A:rtTA); VectorBuilder). Co-expression of WNK1 and GFP is controlled by the doxycycline-inducible TRE promoter. Doxycycline facilitates binding of the rtTA transcriptional enhancer and prevents binding of the tTS transcriptional silencer, thus activating gene expression. A GFP-encoding construct was used as a control to assess transfection efficiency.

### Transfection procedures

Hippocampal neurons were transfected between days in vitro (DIV) 13 and 14 using TransFectin (Bio-Rad), following the manufacturer’s instructions. Briefly, the appropriate plasmid combinations (see Table 1) were mixed in complete culture medium to achieve a total plasmid mass of 1 μg per 20-mm well. Then, TransFectin was mixed with the complete medium at a ratio of 3 μl of TransFectin per 1 μg of plasmid. The plasmid- and TransFectin-containing media were then combined and added to the neurons for 45 minutes. Subsequently, they were replaced with a mixture of conditioned (2/3) and fresh (1/3) complete culture medium. The transfected neurons were used for immunocytochemistry or electrophysiological analysis seven to ten days post-transfection.

**Table 1.** Plasmids ratio used for transfection.

| Experiment | Plasmids ratio for each condition (C) |
| --- | --- |
| Expression and relative membrane expression of recombinant KCC2a- and KCC2b-Flag | C1: KCC2a-Flag (0.8) + EGFP (0.2)<br>C2: KCC2b-Flag (0.8) + EGFP (0.2) |
| Efficacy and specificity of KCC2a-specific shRNAs | C1: shKCC2a_1-GFP (0.2) + KCC2a-Flag (0.8)<br>C2: shKCC2a_3-GFP (0.2) + KCC2a-Flag (0.8)<br>C3: shNT-GFP (0.2) + KCC2a-Flag (0.8)<br>C4: shKCC2a_1-GFP (0.1) + shKCC2a_3-GFP (0.1) + KCC2a-Flag (0.8) |
| Effect of KCC2a knockdown on endogenous KCC2 expression | C1: shKCC2a_1-GFP (0.5) + shKCC2a_3-GFP (0.5)<br>C2: shNT-GFP (1) |
| Effect of KCC2a knockdown on recombinant KCC2b expression and membrane expression | C1: shKCC2a_1-GFP (0.1) + shKCC2a_3-GFP (0.1) + KCC2b-flag (0.8)<br>C2: shNT-GFP (0.2) + KCC2b-flag (0.8) |
| Effect of WNK1 activation on recombinant KCC2b membrane expression (control) | C1: WNK1-CA-GFP (0.5) + KCC2b-Flag (0.5)<br>C2: WNK1-CA-GFP (0.45) + KCC2b-Flag (0.45) + EGFP (0.1) |
| Effect of WNK1 activation on recombinant KCC2b membrane expression (KCC2a knockdown) | C1: WNK1-CA (0.4) + KCC2b-Flag (0.4) + shNT (0.2)<br>C2: WNK1-CA (0.4) + KCC2b-Flag (0.4) + shKCC2a_1-GFP (0.1) + shKCC2a_3-GFP (0.1) |

HEK-293 cells at 70% confluence were transfected using polyethyleneimine (PEI, Sigma-Aldrich) following a protocol similar to that for neuronal transfection. The cells were incubated in the plasmid/PEI mixture for 48 hours prior to fixation.

### Pharmacological treatments

Clathrin-mediated endocytosis was inhibited by incubating neurons for 40 min with the Dynamin Inhibitory Peptide, myristoylated (DiP) (50 μM, MedChemExpress) in complete culture medium at 37°C. Because the effects of DiP are reversible, it was maintained in the medium for all subsequent live staining procedures (20 min). The ubiquitin-proteasome system was blocked by incubating neurons for 2 hr at 37°C with either the ubiquitin-activating enzyme (E1) inhibitor PYR-41 (50 μM, MedChemExpress) or the proteasome inhibitor MG132 (2.5 μM, MedChemExpress) in complete culture medium. As MG132 is a known calpain inhibitor (IC_50_=1.2 μM), this treatment condition also served to inhibit calpain-dependent degradation of KCC2. Expression of the constitutively active WNK1 (WNK1-CA) via the Tet-On system was induced by incubating neurons for 24 hr with doxycycline hyclate (2 μg/ml, Sigma-Aldrich) in complete culture medium at 37°C.

### Immunocytochemistry

To ensure the specificity of the in-house rabbit anti-KCC2a antibody (Markkanen *et al*, 2014), antibodies were pre-adsorbed on brainstems from P9 KCC2a-KO mice. Tissue was fresh-frozen, fixed in 4% PFA overnight at 4°C, and sectioned at 50 μm. Approximately forty sections were permeabilized with 0.25% Triton X−100 (Carl Roth) and incubated for 3 days at 4°C with 1 ml of the rabbit anti-KCC2a primary antibody (1:750) in 3% inactivated goat serum under agitation. The resulting collected antibody solution was subsequently used for KCC2a immunocytochemistry experiments on cultured neurons.

To quantify total endogenous KCC2a and KCC2 expression, the cells were fixed in a solution of 4% paraformaldehyde and 4% sucrose for 15 minutes. The neurons were then permeabilized with 0.25% Triton X-100 and blocked for 30 minutes in 10% inactivated goat serum (Gibco). A two-step primary/secondary antibody staining protocol was then used sequentially. First, the coverslips were incubated with rabbit anti-KCC2a primary antibody (dilution as in Table 2) in 3% inactivated goat serum for one hour at room temperature. After washing with 1x PBS, the cells were incubated with a Cy3-conjugated donkey anti-rabbit secondary antibody solution for 45 minutes at room temperature. The process was then repeated using a rabbit anti-KCC2 antibody and a A647-conjugated donkey anti-rabbit secondary antibody.

**Table 2.**
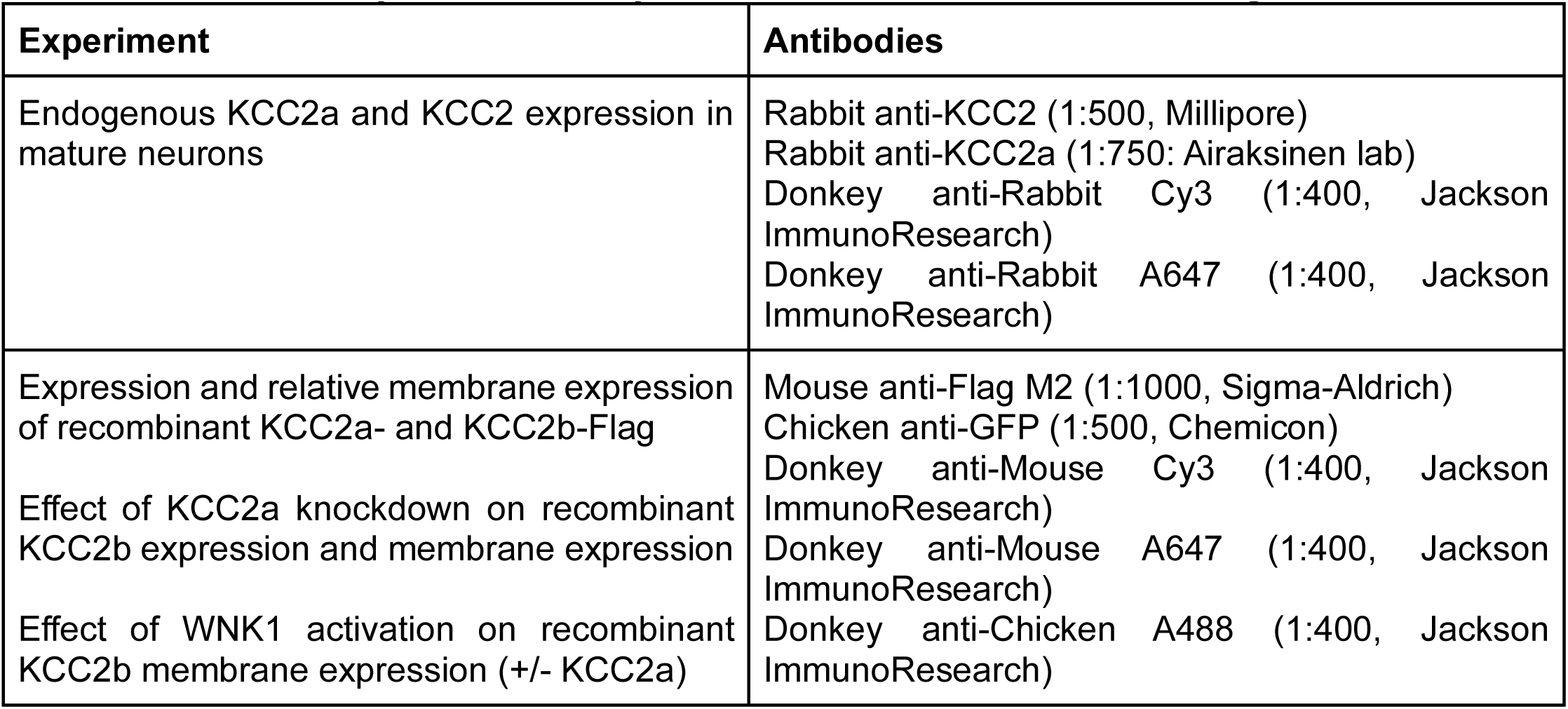
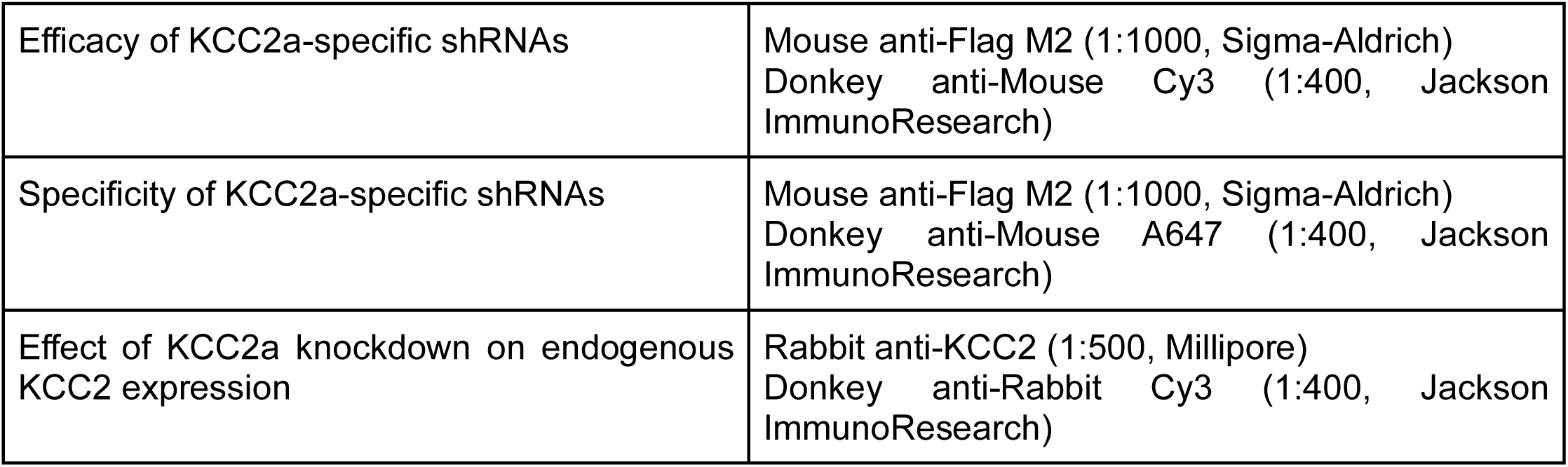
Primary and secondary antibodies used for immunostainings.

**Table 3.**
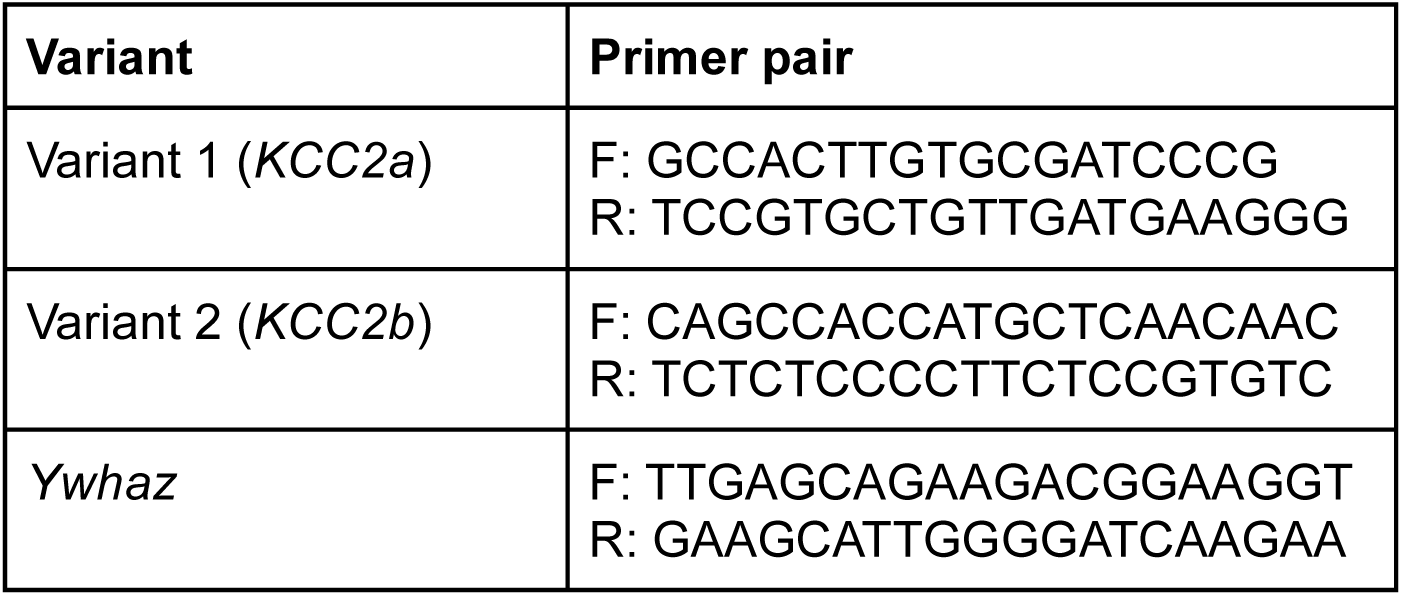
Primers used for the RT-qPCR.

To quantify the relative membrane expression of recombinant KCC2 isoforms, a live-immunostaining protocol was used as previously described (Al Awabdh *et al*, 2022). Living neurons were incubated with mouse anti-flag M2 primary antibodies in MEM medium (Gibco) supplemented with D-Glucose (33 mM), sodium pyruvate (1 mM), HEPES (20 mM), glutamine (2 mM), and B27 (1X) to tag the membrane-inserted KCC2-Flag protein (surface) at 13°C for 20 min. The cells were then fixed, permeabilized and blocked as previously described. After washing with 1x PBS, the cells were sequentially incubated with: a Cy3-conjugated donkey anti-mouse antibody, a mouse anti-Flag M2 and chicken anti-GFP antibody solution and finally a solution containing A647-conjugated donkey anti-mouse and A488 donkey anti-chicken secondary antibodies. The cells were then washed, DAPI-stained, and mounted. For experiments quantifying total endogenous KCC2 or recombinant Flag-tagged KCC2 isoforms, the cells were fixed, permeabilized, and blocked as described above. Primary antibodies (combination as in Table 2) were incubated for one hour at room temperature in 3% inactivated goat serum. After washing with 1X PBS, the cells were incubated with the appropriate secondary antibody solutions (Table 2) for 45 minutes at room temperature. The cells were then washed, DAPI-stained, and mounted.

### Image acquisition and analysis

Standard fluorescence quantification images were acquired using a Leica DM6000 epifluorescence microscope with a 63x objective (NA 1.40) and a 12-bit cooled CCD camera (Micromax, Roper Scientific). The microscope was controlled using Metamorph software (Roper Scientific). The exposure time and other acquisition parameters were determined for each fluorescent channel using the brightest cells to prevent saturation, and these parameters were kept constant across all cells within a given experimental replicate. ImageJ software was used to quantify the surface-to-total fluorescence ratio of recombinant KCC2 isoforms. A region of interest (ROI) was drawn around a primary dendrite positioned in the focal plane. Mean fluorescence intensity was then measured sequentially for the surface (Cy3) and total (Alexa647) pools. Background mean intensity was measured separately for each fluorophore and subtracted from the corresponding signal intensity within the ROI to yield the corrected fluorescence intensity.

Image acquisition for experiments including endogenous KCC2a expression and KCC2 expression depending on various treatments was performed using a Leica SP5 confocal microscope with a 63x objective (NA 1.40) and LAS AF software. Confocal images were acquired at a zoom level of 5. For each neuron, a Z-stack was acquired across a primary dendrite with 0.5 μm Z-steps. For analysis, the Z-stacks were processed into a sum projection using ImageJ. A primary dendrite was selected, and an ROI was drawn. Background fluorescence was subtracted from the A647 signal to determine total protein expression. To account for dendrite thickness, the value was normalized by the dendrite thickness, which was calculated as [number of images in stack − 1] × 0.5 μm. Relative surface expression was calculated as the ratio of corrected surface intensity to corrected total intensity.

We analyzed panKCC2 and KCC2a expression along the somato-dendritic axis using Z-axis projected images. The number of planes was adjusted depending on whether somatic or dendritic expression was quantified. Background intensity was measured independently for the soma stack and the dendrite stack to ensure accurate subtraction. Normalization by the thickness of the neuron ([number of images in the stack – 1]*0.5) was performed independently at the soma and at the dendrite.

IMARIS software was used to quantify enrichment of panKCC2 and KCC2a in protein clusters. Dendritic segments were visualized in 3D and isolated from background noise. Aggregates with a diameter greater than 0.7 μm and a fluorescence intensity exceeding a user-defined threshold were classified as clusters and modeled as 1-μm spheres. The number of clusters and the fluorescence intensity within the clusters and within the dendritic portion were measured for both isoforms, and the corresponding background values were subtracted.

Analysis of KCC2 and recombinant KCC2b-Flag clusters was performed using an in-house Metamorph routine as previously described (Al Awabdh *et al*., 2022). A primary dendrite ROI was selected, and images were digitally flattened and filtered. Clusters were detected using a user-defined intensity threshold and a minimum area of 0.031 µm^2^. The cluster regions were superimposed on the raw images to calculate their mean density, area, and integrated intensity.

### RNA extraction

RNA was extracted from rat brain tissue using TRIzol reagent (Invitrogen). The tissue was snap-frozen on dry ice and stored at −80°C. The tissue was lysed by immersing it in 1 ml of TRIzol reagent and homogenizing it with a Potter homogenizer. Then, 200 μl of chloroform was added, and the mixture was centrifuged to separate the aqueous phase containing the RNA. The RNA was then precipitated by adding 500 μl of isopropanol, followed by centrifugation for 10 minutes. The RNA pellet was washed with 75% ethanol, air-dried for 10 minutes, and resuspended in 50 μl of RNase-free water.

RNA was extracted from cultured neurons using the RNAqueous-Micro Kit (Ambion). Neurons were plated in 6-well plates at a density of 3.5×10⁵ cells/well. For each time point (DIV 7, 10, and 24), neurons from two wells were harvested and pooled. The cells were detached by mechanical dissociation and centrifuged. The supernatant was discarded, cells were lysed, and 50 μl of absolute ethanol was added to the lysate. The lysate/ethanol mixture was loaded onto a column provided in the kit, followed by a series of washing and centrifugation steps. RNA was eluted in 20 μl elution buffer. Residual genomic DNA was removed by DNase I treatment.

### Reverse transcription and quantitative PCR

Total RNA was reverse transcribed using SuperScript II reverse transcriptase (Invitrogen). The reaction mixture contained 1 μg RNA, 100 ng random primers (Thermo Fisher Scientific), and 1 μl of a 10 mM dNTP mix (Thermo Fisher Scientific) and was brought to 13 μl with sterile water. The mixture was denatured at 65°C for 5 minutes. DTT was added, followed by a 2 min incubation at RT, and then 1 μl SuperScript reverse transcriptase was added. The synthesis steps were 10 minutes at 25°C and 50 minutes at 42°C. The reaction was then inactivated with a 15-minute incubation at 70°C.

Quantitative PCR was conducted using the ABsolute Blue QPCR Mix, SYBR Green, ROX (Thermo Fisher Scientific). Each target amplification (KCC2a, KCC2b, and the housekeeping gene Ywhaz) was run in triplicate in a 384-well qPCR plate, alongside a no-reverse-transcriptase (NRT) control and a water control. Each 10 μl reaction consisted of: 5 μl Tp2X Sybr, 4 μl template cDNA (C=10 ng/μl), 0.2 μl forward and reverse primer (10 μM), and 0.8 μl water. qPCR was performed on a QuantStudio5 device using the following thermal cycling program: initial denaturation at 95°C for 10 min, followed by 40 cycles of 95°C for 30 s, 58°C for 30 s, 72°C for 20 s, and a final melt curve step at 95°C for 15 s.

The RT-qPCR data were analyzed as follow. For each condition (rat hippocampus/cortex, hippocampal neurons in culture), 3-4 independent experiments were performed. Within a single experiment, each variant (*KCC2a*, *KCC2b*, *Ywhaz*) was analyzed at different times (E18/adult or DIV7/10/24) and each association of variant and time was run in triplicate. We calculated the mean Ct of the triplicates for each variant-time, and the value of 2^-ΔΔCt^ was calculated by subtraction of a variant mean Ct by *Ywhaz* mean Ct and then by normalization either by KCC2b value or by the E18/DIV7 value. Finally, we calculated the average 2^-ΔΔCt^ of the independent experiments for each variant-time.

### Western blot

Brains used for the western blots were obtained by fast cervical dislocation and decapitation. They were then dissected out, snap frozen by immersion in isopentane and stored at -80°C. Adult brain hemispheres or P9 whole brains (with brainstem dissected out) from KCC2a-KO C57BL6/J mice and WT littermates were lysed in a modified RIPA buffer containing (in mM): 50 Tris-HCl (pH 7.4), 150 NaCl, 1% Nonidet P−40, 1% Triton X-100, 0.5% DOC, 0.1% SDS, 50 NaF, 1 EDTA supplemented with 1X protease inhibitors (Roche) and phosphatase inhibitors (Thermo Fisher Scientific). The lysis volume was standardized (10 μl RIPA per 1 mg tissue). Lysis was achieved by homogenization in a Potter homogenizer. Samples were centrifuged, and the supernatant containing the protein lysate was collected. Protein concentration was determined using the Pierce^TM^ BCA Protein assay (Thermo Fisher Scientific).

For KCC2 and SPAK western blots, protein extracts were prepared with sample reducing agent (Invitrogen) and LDS sample buffer (Invitrogen) to a final concentration of 3 μg/μl and denatured for 1 hr at 37°C. 10 μl of sample was loaded onto a 4-12% Bis-Tris polyacrylamide gel (Invitrogen) for electrophoresis. Proteins were transferred to a nitrocellulose membrane for 3 hr at 100 V at 4°C. Membranes were washed (3×10 min) with TBS-Tween (TBST) and blocked with 5% (w/v) skim milk. Primary antibodies were incubated overnight at 4°C in 5% (w/v) skim milk solution: rabbit anti-KCC2 (1:1000, Upstate), rabbit anti-KCC2pT1007 (1:1000, PhosphoSolutions), sheep anti-SPAK (1:50, MRC PPU reagents Dundee), sheep anti-SPAKpS373 (1:50, MRC PPU reagents Dundee) and rabbit anti-GAPDH (1:2000, Invitrogen) as loading control. Membranes were washed (3×10 min TBST) and incubated for 1 hr at RT with goat anti-rabbit DyLight 800 secondary antibody (1:5000, LI-COR) and donkey anti-sheep 800 secondary antibody (1:1000, Rockland, for SPAK staining) in 1% (w/v) skim milk. Fluorescence was acquired using the Odyssey infrared imaging system (LI-COR Bioscience).

For WNK1 western blots, protein samples were denatured for 10 min at 70°C. 10 μl of protein was loaded on a 3-8% Tris-Acetate polyacrylamide gel (Invitrogen) and transferred to a nitrocellulose membrane for 2.5 hr at 65 V at 4°C. Immunostaining utilized sheep anti-WNK1 (1:50, MRC PPU reagents Dundee) or sheep anti-WNK1pS382 (1:50, MRC PPU reagents Dundee) primary antibodies, followed by donkey anti-sheep 800 secondary antibody (1:1000, Rockland). Integrated intensity of the different bands signal was measured using ImageJ software. To quantify KCC2 (or SPAK) expression for each animal, the integrated intensity of the KCC2 signal was divided by the integrated intensity of the corresponding GAPDH signal. To quantify KCC2 phosphorylation level for each animal, the integrated intensity of the KCC2 Thr1007 phosphorylated signal was divided by the corresponding GAPDH (1) signal. The integrated intensity of the KCC2 signal was divided by the corresponding GAPDH (2) signal. Finally, the 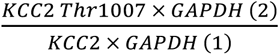 ratio was calculated. To quantify WNK1 phosphorylation level for each animal, the integrated intensity of the WNK1 Ser382 phosphorylated signal was divided by the integrated intensity of the corresponding WNK1 signal. To account for technical variability, the final protein/GAPDH or phospho-protein/protein ratio for each biological sample (animal) was averaged from three independent technical replicates.

### Electrophysiology

Whole-cell patch clamp recordings were performed on DIV 21-24 hippocampal neurons superfused at a rate of 2 ml/min with an artificial cerebral fluid solution (ACSF) composed of (in mM): 2 CaCl_2_, 2 KCl, 3 MgCl_2_, 10 HEPES, 20 D-glucose, 120 NaCl at 34°C. Neurons were recorded using 5 MOhm borosilicate glass pipettes filled with (in mM): 104 K-gluconate, 25.4 KCl, 10 HEPES, 10 EGTA, 2 MgATP, 0.4 Na_3_GTP, 1.8 MgCl_2_. The liquid junction potential (14.9 mV) was corrected *post hoc*. Signals were acquired with a Multiclamp 700B amplifier (Molecular Devices), low-pass filtered at 2 kHz and digitized at 20 kHz using Clampex 11 (Molecular Devices). GABA_A_R-mediated currents were isolated using (in µM): 1 TTX (Na^+^ channel blocker, Latoxan), 50 D-AP5 (NMDA receptor antagonist, HelloBio), 10 NBQX (AMPA receptor antagonist, HelloBio) and 100 CGP54626 (GABA_B_ receptor antagonist, Tocris Bioscience). After achieving whole-cell configuration, an isoguvacine-filled pipet (100 μM) was positioned near the soma. Voltage steps ranging from −80 mV to 0 mV (10 mV step) were applied, and GABAAR-mediated currents were evoked by the release of isoguvacine at each step. The isoguvacine pipet was then repositioned over a primary dendrite at a distance of 50−100 μm from the soma, and the same voltage-step protocol coupled with isoguvacine release was applied. Currents were registered at each holding potential, corrected for both the access resistance voltage drop and the liquid junction potential. Current-voltage (I-V) curves were plotted offline using Clampfit 11 to estimate the reversal potential of GABA_A_R currents at the soma (EGABA_soma_) and the dendrite (EGABA_dendrite_). The difference (ΔE_GABA_=EGABA_dendrite_−EGABA_soma_) was computed and then normalized by the distance between the two application sites to quantify the EGABA gradient.

### Statistical analysis

All statistical procedures were conducted in SigmaPlot 13, with significance set at p<0.05. While data are visualized as box plots showing the median and quartiles, numerical differences discussed in the text are based on mean values. We determined the appropriate test by first evaluating normality (Shapiro-Wilk) and, for multi-group comparisons, equal variance (Brown-Forsythe). Two-group comparisons were conducted using two-tailed Student’s t-test (normal distribution) or Mann–Whitney rank sum test (non-normal). For specific experiments involving more than two groups, such as pharmacological rescue experiments (e.g., Fig. 5), p-values were further corrected using the Holm-Šidák method. Multi-group comparisons were performed using one-way ANOVA with Tukey’s post hoc test, or Kruskal–Wallis ANOVA on ranks with Dunn’s post hoc test if assumptions were violated.

## Acknowledgements

The authors wish to thank Pavel Uvarov for advice, technical help and critical reading of the manuscript, and Muriel Thoby-Brisson for providing brain samples from KCC2a KO and WT mice. P.W. was the recipient of a scholarship from the Erasmus Mundus program. This work was supported in part by ANR (ANR-21-NEU2-0007-02 and ANR-21-CE16-0006-03), Inserm and the Paris Brain Institute. We are also grateful to the Imaging Facility of the Institut du Fer à Moulin for assistance with confocal imaging and data analysis. The Poncer lab is affiliated to DIM C-BRAINS, supported by the Conseil Régional d’Île-de-France.

## Authors contribution

Conceptualization: CP, JCP, PW; Formal Analysis: CP, PW; Funding acquisition: JCP; Investigation: CP, PW; Methodology: CP, PW, JCP; Resources: MR, MM, MA; Supervision: JCP; Visualization: CP, JCP; Writing – original draft: CP, JCP; Writing – review & editing: CP, JCP, MM, MA.

